# Exosomes are key regulators of non-cell autonomous communication in senescence

**DOI:** 10.1101/356238

**Authors:** Michela Borghesan, Juan Fafián-Labora, Paula Carpintero-Fernández, Pilar Ximenez-Embun, Hector Peinado, Javier Muñoz, Ana O’Loghlen

## Abstract

Senescence is a cellular phenotype characterized by an irreversible cell cycle arrest and the secretion of inflammatory proteins, denominated senescence-associated secretory phenotype (SASP). The SASP is important in influencing the behavior of neighboring cells and altering the microenvironment; yet, until now this role has been mainly attributed to soluble factors. Here, we report that extracellular vesicles also alter the environment by transmitting the senescent phenotype to other cells via exosomes (extracellular vesicles of endocytic origin). A combination of functional assays, Cre-/oxP reporter systems, proteomic analysis and RNAi screens confirm that exosomes form part of the senescent secretome and mediate paracrine senescence via the activation of a non-canonical interferon (IFN) pathway. Altogether, we speculate that exosomes could be drivers of tissue degeneration both locally and systemically during aging and age-related disease.

## INTRODUCTION

The establishment of cellular senescence is categorized by an irreversible cell cycle arrest and the capacity to modify the microenvironment through a particular secretome named SASP (senescence-associated secretory phenotype). The activation of senescence is a response to different stress triggers in order to prevent the propagation of damaged and has been show to occur *in vitro* and *in vivo.* In fact, senescence has been observed *in vivo* during both biological and pathological processes such as development, cancer, fibrosis and wound healing (He and Sharpless, 2017; Munoz-Espin and Serrano, 2014). The SASP controls its surroundings by reinforcing senescence in an autocrine (cell autonomous) and paracrine (non-cell autonomous) manner, by recruiting immune cells to eliminate senescent cells and by inducing a stem-cell like phenotype in damaged cells (Mosteiro et al., 2016; Salama et al., 2014). Altogether, the SASP provides the necessary balance to restore tissue homeostasis when it has been compromised. Paradoxically, the SASP can also contribute to enhance tissue damage, but the mechanisms behind this pleiotropic activity of the SASP are not well understood.

Most studies *in vitro* and *in vivo* have attributed the diverse functions of the SASP to individual protein components such as IL-6 or IL-8 to reinforce autocrine senescence (Acosta et al., 2008; Kuilman et al., 2008) or TGF-β as the main mediator of paracrine senescence (Acosta et al., 2013; Rapisarda et al., 2017) or to a dynamic SASP with TGF-β or IL-6 as predominant individual components (Hoare et al., 2016). However, it is still unclear how these diverse SASP components regulate senescence. In fact, inhibition of the SASP by block the mammalian target of rapamycin (mTOR) only partially blocks the SASP, suggesting that alternative mechanisms might exist (Herranz et al., 2015; Laberge et al., 2015).

Exosomes are small extracellular vesicles (30-120 nm) of endocytic origin secreted by all cell types and therefore found in most bodily fluids. Exosomes contain nucleic acids, proteins and lipids that generally reflect the status of the parental cell and can influence the behavior of recipient cells locally and systemically (Tkach and Thery, 2016). In fact, the increasing literature regarding exosomes show that they can be used as disease biomarkers (Melo et al., 2015), indicators of cancer metastatic predisposition (Hoshino et al., 2015) or as therapeutic carrier vesicles (Kamerkar et al., 2017). However, although some studies have found an increase in the number of exosomes released during senescence (Lehmann et al., 2008; Takasugi et al., 2017), very little is known regarding the role that exosomes play as SASP mediators in the microenvironment.

Here, our data show that inhibition of exosome biogenesis prevents the induction of senescence in neighboring cells and that both the soluble and exosome fraction are capable of transmitting senescence. Furthermore, the analysis of the individual cells internalizing exosomes, show a direct link between the uptake of exosomes derived from senescent cells and paracrine senescence induction. We also show that exosome biogenesis is an essential cellular process. Interestingly, exosome protein characterization by Mass Spectrometry (MS) followed by a functional siRNA screen identify the non-classical interferon (IFN) pathway as responsible for transmitting senescence to normal cells.

## RESULTS

### Blocking exosome biogenesis prevents the paracrine induction of senescence

In order to investigate whether exosomes act as intercellular mediators in neighboring cells during senescence, we took advantage of HFFF2 human foreskin primary fibroblasts expressing a chimeric fusion protein (ER) with either an empty vector (ER:EV) or expressing the oncogene H-RAS^G12V^ (ER:RAS) (Rapisarda et al., 2017), termed iC or iRAS respectively. We blocked exosome biogenesis in iRAS treated with 200nM 4- hydroxytamoxifen (4OHT) using either small molecules inhibitors or siRNA targeting RNAs/proteins essential for exosome biogenesis. We next washed the treated cells, to avoid the carry-over of siRNA/inhibitors, added fresh media for 72h and treated normal HFFF2 for an additional 72h with the different conditioned media (CM) (Figure 1A-C). As previously described, the CM from iRAS cells (mimicking oncogene-induced senescence, OIS) induced cell cycle arrest in normal HFFF2s, measured by determining the percentage of cells incorporating BrdU. However, HFFF2s incubated with the CM derived from iRAS treated with increasing concentrations of two different small molecule inhibitors, GW4869 and Spiroepoxide (SpE) that block the enzyme neutral sphingomyelinase (N-SMase), which regulates exosome biogenesis and release (**Figure S1A**) (Hannun and Obeid, 2008; Trajkovic et al., 2008), prevents this proliferation arrest (Figure 1B-**left graph**). Torin-2, an mTOR inhibitor previously described to suppress the SASP (Herranz et al., 2015; Laberge et al., 2015) also prevented paracrine senescence (Figure 1B). It has been described that blocking exosome biogenesis induces a DNA-damage response (Takahashi et al., 2017), so next we decided to determine the levels of phosphorylated-γH2AX (p-γH2AX), which is also a marker of senescence, by immunofluorescence (IF). To our surprise, the levels of p-γH2AX also decreased when both GW4869 and SpE were used and also upon the use of Torin-2 (Figure 1B-**right graph**). To further confirm the implication of exosomes on transmitting senescence, we used a non-targeting small interfering RNA (siRNA) (Scr) and two siRNA targeting two canonical regulators of exosome secretion, RAB27A (siRAB27A #3,#4) (Ostrowski et al., 2010) and Synaptotagmin-7 (siSYT7 #1,#2) (Hoshino et al., 2013) (**Figure S1B,C**). Our data show that iRAS transfected with siRAB27A and siSYT7 prevented exosome release (**Figure S1C)** and therefore their CM did not induce growth arrest nor DNA-damage (Figure 1C). We also analyzed the expression levels of additional markers of senescence by IF such as p21^CIP^ (**Figure S1D,E**) and p53 (not shown). A similar response was observed in an additional strain of fibroblasts, IMR-90, by measuring the levels of expression of p21^CIP^ by IF (**Figure S1F**) and p-γH2AX (not shown). Next, we used Transwell inserts with 0.4 μ.m pore membranes to avoid the transfer of larger vesicles, plating iC and iRAS with the different treatments in the upper chamber (UC) and later plating HFFF2 in the lower chamber (LC) with fresh CM (Figure 1D). As per our previous data, we could observe that treatment with GW4869 (10μM) and SpE (5μ,M) prevented paracrine senescence in a similar fashion to Torin-2 (50nM), observed by a reduction in the cells incorporating BrdU and the expression levels of p16^INK4A^ (Figure 1E, **Figure S1G**). Altogether, our data suggest that exosome release could be implicated in inducing paracrine senescence.

**Figure 1.**
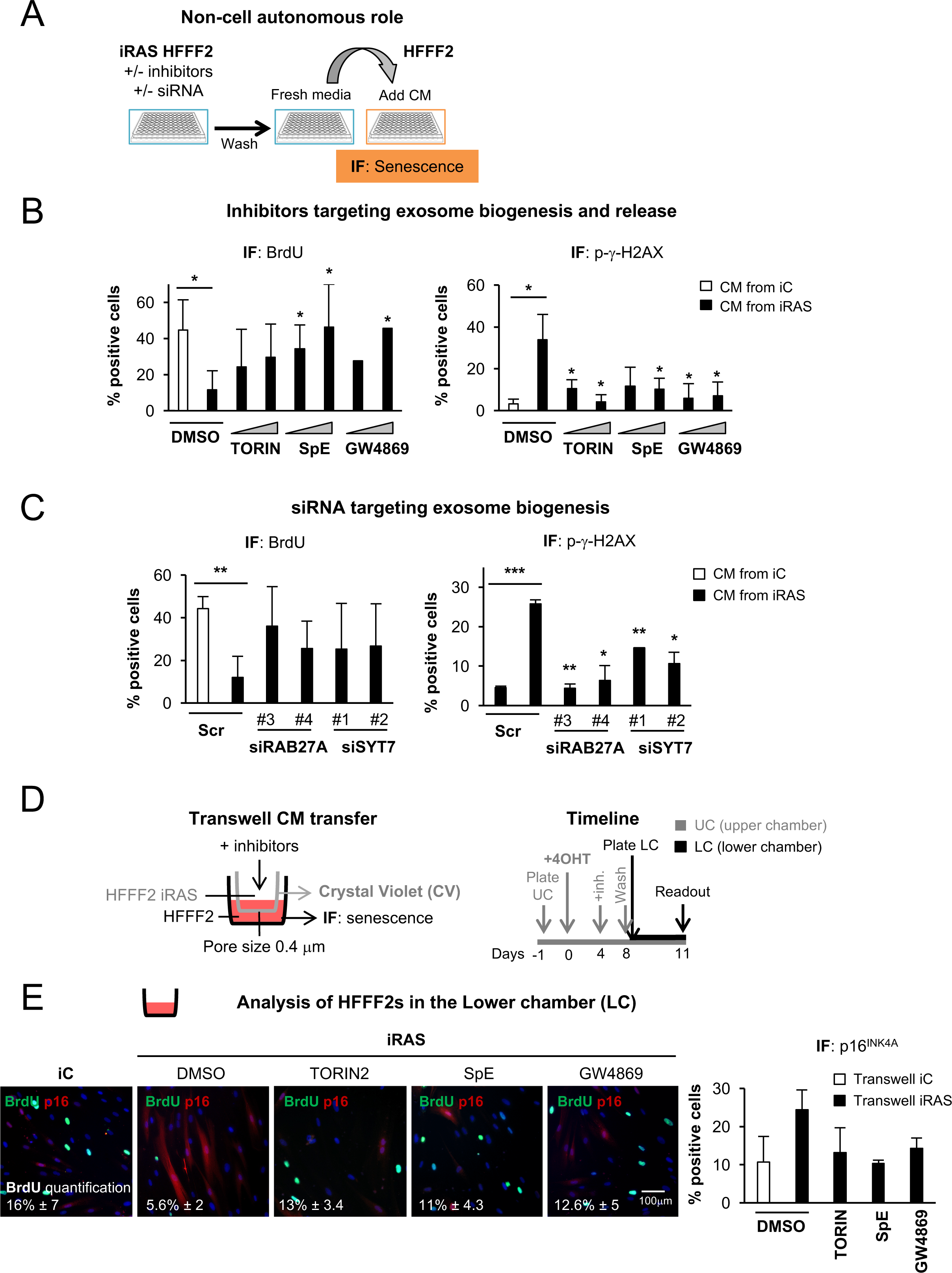
Blocking exosome biogenesis prevents paracrine senescence. (A) Schematic representation of the experimental settings to determine if exosomes mediate paracrine senescence. Briefly, HFFF2 human primary fibroblasts expressing a vector encoding an inducible form of H-RAS^G12V^ ER:RAS (iRAS) were treated with 200nM 4OHT for 2 days followed by siRNA or inhibitors affecting exosome biogenesis. HFFF2 fibroblasts expressing an empty vector (iC) were treated with 200nM 4OHT and used as a control. After the incubation with the inhibitors or transfection with the siRNA, cells were washed and allowed to produce Conditioned Media (CM) for 72h. Normal HFFF2 were then incubated with this CM for further 72h. (B) iRAS HFFFs were treated with 4OHT for 2 days followed by treatment with different concentrations of Torin-2 (25, 50 nM; blocking the release of soluble factors) or two independent inhibitors that block exosome biogenesis: GW4869 (1, 1μM) and spiroepoxide (SpE) (2, 5μM) for 3 days. After treatment, the cells were washed and allowed to produce fresh CM for 72h. CM- treated HFFF2 fibroblasts were then stained to assess for the percentage of cells expressing markers of senescence: incorporation of BrdU and phospho-γH2AX (p- γH2AX) by IF (mean ± SD of 3-4 independent experiments). (C) iRAS HFFF2s were treated with 4OHT for 2 days and then transfected with a scramble siRNA (Scr) or two independent siRNAs targeting RAB27A (#3, #4) or synaptogamin-7 (SYT7 #1, #2) for 3-5 days. iC and iRAS cells were washed and allowed to produce CM free of siRNA for a further 72h. The CM was used to treat HFFF2 for 72h after which the percentage of cells staining positive for BrdU and p-γH2AX was measured and quantified. The graph shows the percentage of cells staining positive for BrdU and p-γH2AX (mean ± SD of 2-3 independent experiments). (D) Schematic representation of the experimental settings and timings to test the implication of exosomes using the Transwell system with a membrane pore size of 0.4μm. In brief, iRAS cells were plated in the upper chamber (UC) and treated with 200nM 4OHT for 2 days and either 50nM Torin-2, 10μM GW4869 or 5μM SpE for 3 further days. At this point HFFF2s were plated in the lower chamber (LC) and left in contact with the upper chamber for 72h. (E) The lower chamber was stained to quantify the percentage of cells incorporating BrdU and expressing p16^INK4A^ by IF. Representative pictures and the quantification for BrdU incorporation are shown on the left panel and the quantification for p16^INK4A^ on the right. Data shows the mean ± SD of 2-3 independent experiments. Scale bar, 100μm. **See also Figure S1.**

### Exosomes derived from iRAS cells are responsible for mediating paracrine senescence

In order to confirm that exosomes are responsible for inducing paracrine senescence we decided to dissect the CM from iC and iRAS HFFF2s and compared the role of large (microvesicles; MV) and small (exosomes; Ex) extracellular vesicles in inducing paracrine senescence on normal HFFF2s (Figure 2A). The treatment of HFFF2s with the whole CM from iC and iRAS show that senescence can be transmitted as seen by a decrease in cell proliferation (staining with crystal violet) and an increase in the percentage of cells staining positive for SA-β-Gal (**Figure S2A**). We next isolated microvesicles (MV), formed by budding of the plasma membrane and exosomes (Ex) from the same CM following the serial ultracentrifugation protocol (Thery et al., 2006) and compared their effect in inducing senescence with the supernatant fraction (SN) containing the soluble fraction (CM depleted of MV and Ex). The SN and exosome fraction from iRAS cells both induced paracrine senescence in normal HFFF2 compared to iC, as shown by a decrease in BrdU incorporation, an increase in p-γH2AX positive cells and the accumulation of p53 by IF (Figure 2B-C, **Figure S2B**). In addition, an increase in p21^CIP^, p16^INK4A^, integrin avp3, previously described to regulate senescence (Rapisarda et al., 2017), could be observed by immunoblotting in HFFF2s treated with Ex isolated from iRAS (Figure 2D). However, we could not observe differences in growth between HFFF2s treated with MVs isolated from either iC or iRAS, nor DNA damage response (**Figure S2B**) nor accumulation of p53 (not shown). We next confirmed the presence of exosomal proteins (TSG101 and ALIX) in our isolated exosomes by immunoblotting and their morphology and size by transmission electron microscopy (TEM) (**Figure S2C,D**). Furthermore, treatment of HFFF2s with increasing concentrations of exosomes derived from iRAS cells induced a dose-dependent senescent response as shown by quantifying the percentage of cells staining positive for SA-β-Gal and the levels of p53 and BrdU positive cells by IF (**Figure S2E**). Thus, exosomes isolated from iRAS cells are capable of inducing senescence in normal HFFF2.

**Figure 2.**
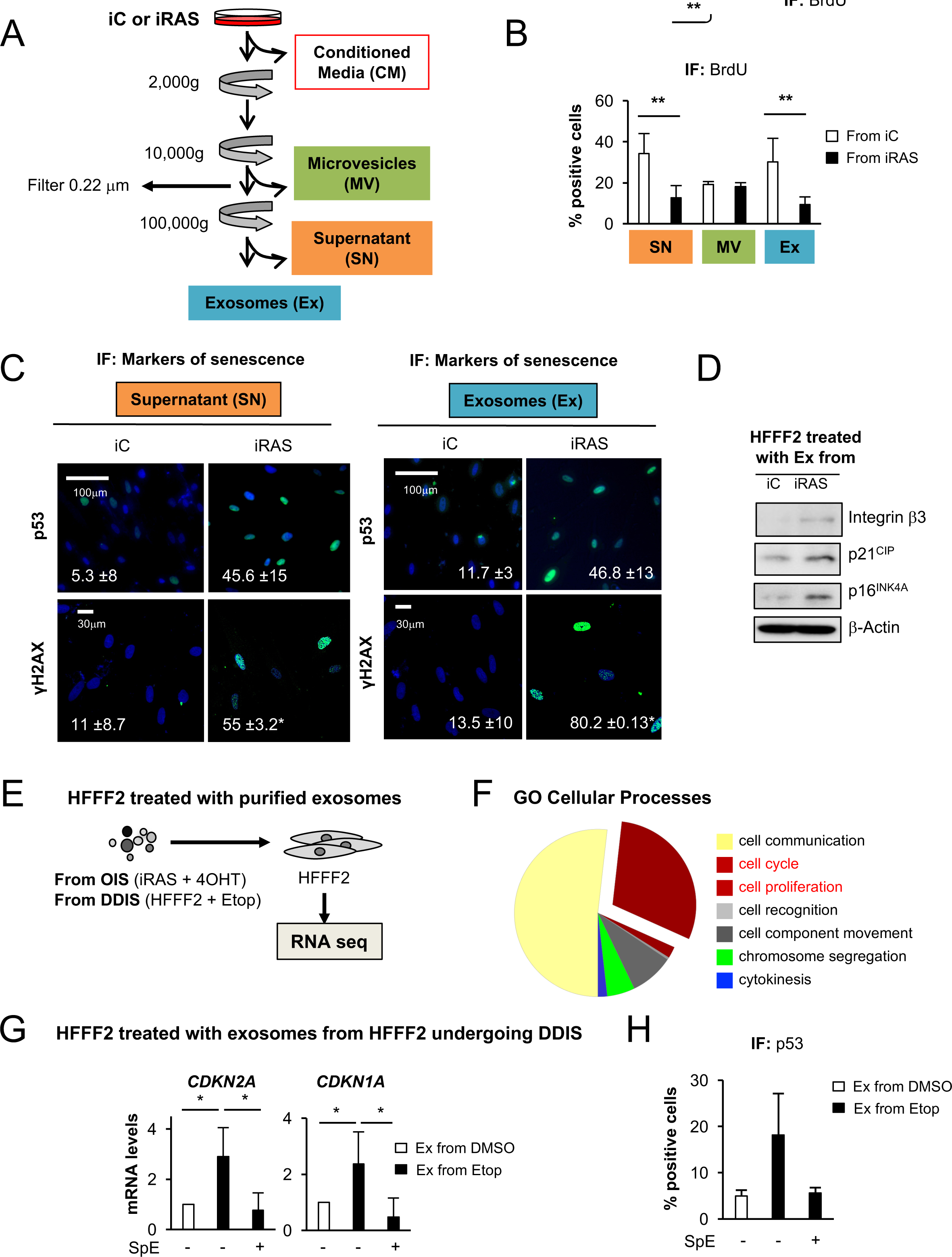
Exosomes form part of the senescent secretome and mediate a cell cycle arrest in normal HFFF2s. (A) Schematic representation of the proof-of-concept experiments performed to show that exosomes form part of the senescent secretome. CM was taken from iC or iRAS HFFF2s and tested for the ability to induce senescence in HFFF2 as a whole (**Figure S2A**) or (B) processes by serial centrifugation to evaluate the effect of the different fractions, supernatant (SN), microvesicles (MV) or exosomes (Ex), to induce senescence in HFFF2s. (B) HFFF2 fibroblasts were treated for 72h with the different fractions of the CM: SN, MV or Ex from iC or iRAS cells and the percentage of cells incorporating BrdU by IF was quantified. The graph represents the mean ± SD of 2-5 independent experiments. (C) Representative pictures show HFFFs treated for 72h with either the SN or the Ex fraction and stained for p53 or p-γH2AX. The quantification of the percentage of HFFF2s staining positive for each antibody is shown (mean ± SD of 2 independent experiments). (D) Immunoblotting for HFFF2s treated for 72h with Ex derived from iC and iRAS show an increase in p21^CIP^, p16^INK4A^ and integrin β3 when treated with iRAS Ex. β- Actin is used as loading control. (E) RNA from HFFF2 cells treated for 72h with exosomes isolated from iRAS (mimicking OIS) or HFFF2 treated with Etop (mimicking DDIS) was isolated from 3 independent experiments and sent for RNA sequencing. (F) Gene ontology (GO) analysis for genes involved in Cellular Processes with > 2 log_2_ fold differential expression and p-value < 0.05 in both OIS- and DDIS-treated HFFF2s. The pie chart shows a high proportion of genes related to the “Cell cycle” and “Cell proliferation” pathways. (G,H) To confirm the RNA sequencing data, HFFF2 were incubated with an equal number of exosomes from HFFF2 treated with Etop with or without SpE and (G) the relative mRNA levels were determined for genes implicated in cell cycle regulation *(CDKN1A, CDKN1A)* by qPCR or (H) p53 protein levels were determined by IF. Data represent mean ± SD of (G) 2-3 and (H) 2 independent experiments. **See also Figure S2.**

### Exosomes isolated from iRAS induce a cell cycle arrest in HFFFs

To identify the pathways regulated upon exosome treatment, we decided to perform RNA sequencing of HFFF2s incubated with exosomes derived from iRAS and from HFFF2 treated with 50μM Etoposide for 48h followed by 5 days incubation with fresh medium (mimicking DNA-damage induced senescence; DDIS) (Figure 2E). Gene ontology (GO) analysis of genes deregulated by > 2 log_2_ fold difference expression levels and p-value < 0.05 in both OIS- and DDIS-exosome treated HFFF2s, confirmed a shared ‘‘Cell cycle” and “Cell proliferation” signature (Figure 2E,F), with the “p53 signaling pathway’ being over-represented (**Figure S2F**). In fact, HFFF2s treated with an equal number of control and DDIS-derived exosomes show an increase in the mRNA expression levels of the cell cycle regulators, *CDKN1A, CDKN2A*, by qPCR and p53 protein levels by IF (Figure 2G,H), while SpE prevented this upregulation (Figure 2G,H). Similar results were obtained in HFFF2 treated with exosomes derived from iRAS cells treated with and without SpE (**Figure S2G**). These data indicate that the inhibition of exosome biogenesis alters the content of exosomes during senescence, suggesting a specific cargo loading exists during senescence independent of exosome number.

### The uptake of exosomes derived from senescent cells induces paracrine senescence

Next, we decided to investigate whether exosomes from both iC and iRAS cells were being internalized by HFFF2. For this we generated iRAS HFFF2 cells expressing a mCherry-CD63 construct (iRAS;CD63-ch), which release mCherry positive exosomes (CD63-ex), and HFFF2s expressing GFP (GFP). First, we decided to co-culture both GFP^+^ and iRAS;CD63-ch^+^ cells in a 1:1 ratio with 200nM 4OHT for 2 days followed by replenishing with fresh media for 4 days (Figure 3A) and determine the presence of CD63-ex transfer in GFP^+^ HFFF2s by confocal microscopy (**Figure S3A**) and the super resolution Airyscan microscope (Figure 3B). In fact, using both techniques we could observe CD63-ex transfer within GFP^+^ HFFF2s (Figure 3B, **Figure S3A**). Furthermore, as shown in Figure 3B, 3D rotation of a Z- stack image shows internalized CD63-ex. In order to avoid confounding effects of the SASP, we next purified CD63-ex from iRAS;CD63-ch and iC;CD63-ch cells and treated normal unlabeled HFFF2 cells (Figure 3C). To determine whether the uptake of CD63-ex from iRAS;CD63-ch was inducing senescence in HFFF2, we next quantified the percentage of cells with CD63-ex uptake (CD63-ex^+^ HFFF2) and determined how many of this selected population of cells were proliferating. This was done by staining with the proliferation marker Ki67 by IF. The quantification of the percentage of CD63-ex^+^/Ki67^+^ cells shows that HFFF2 treated with exosomes purified from iRAS;CD63-ch that have taken up these exosomes (e.g. CD63- ex^+^ HFFF2) proliferate less (less Ki67^+^) (Figure 3C; **right panel**). These data suggest that exosomes derived from iRAS are the cause of paracrine senescence in HFFF2s.

**Figure 3.**
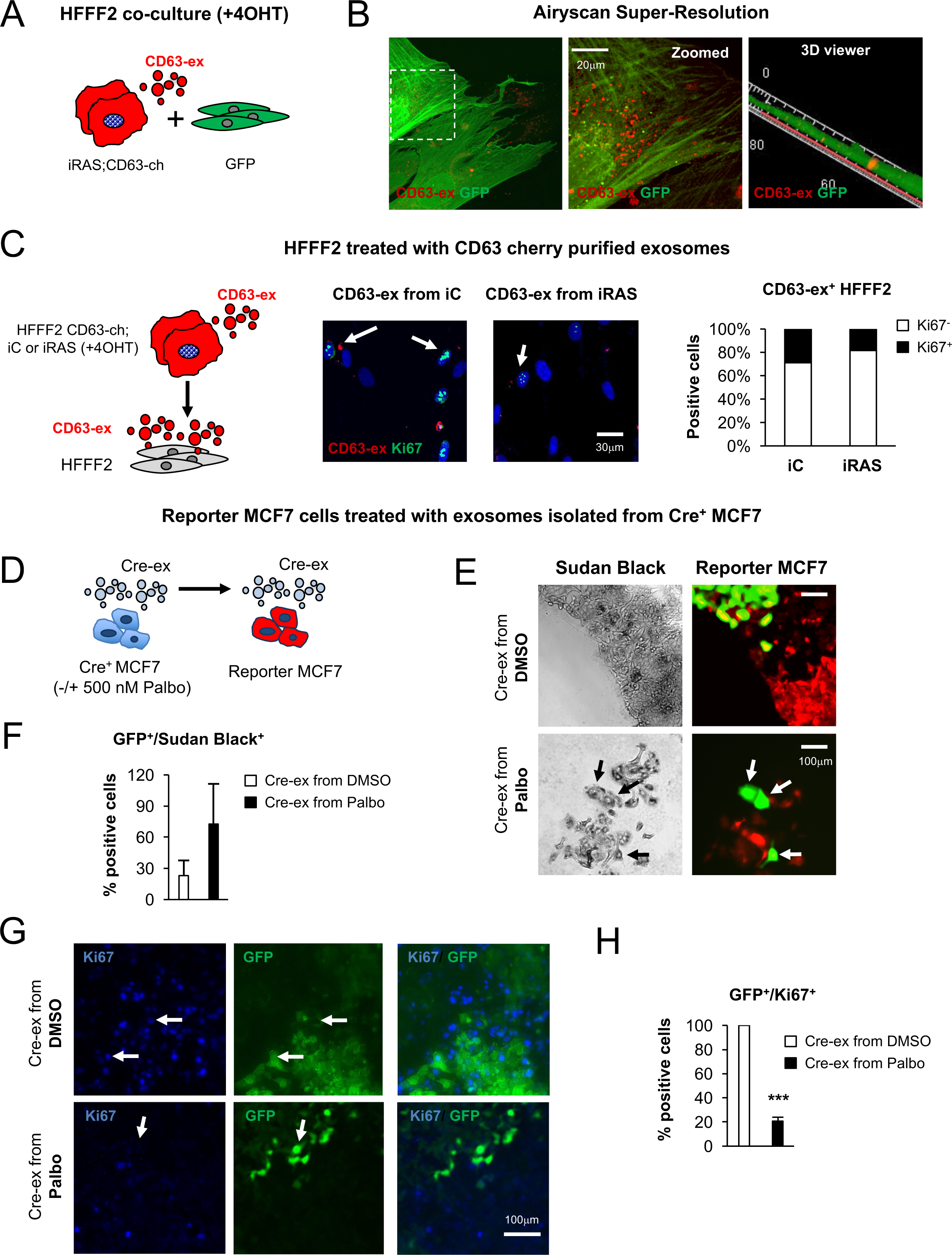
The uptake of exosomes derived from cells undergoing senescence induces paracrine senescence. (A) Schematic representation of HFFF2 fibroblasts used for the co-culture experiments. (A,B) Co-culture of HFFF2 expressing a GFP plasmid (GFP) and iRAS HFFF2 fibroblasts expressing a retroviral construct encoding for mCherry-CD63 (iRAS;CD63-ch). Cells were plated at a 1:1 ratio and treated with 4OHT for 48h followed by 3-4 days with fresh media. (B) Representative images showing the uptake of CD63-cherry positive exosomes (CD63-ex) in GFP cells acquired with the super-resolution microscope Airyscan. Right panel shows a 3D reconstruction of confocal Z-stack images showing CD63-ex inside GFP cells. Scale bar, 20μm. (C) HFFF2 fibroblasts were treated for 72h with exosomes isolated from iRAS;CD63-ch or HFFF2 cells expressing a vector control and mCherry-CD63 (iC;CD63-ch). Representative images show the uptake of CD63-ex in HFFF2s treated with exosomes isolated from both iC and iRAS. The graph shows the quantification of HFFF2s that have successfully internalized CD63-ex (Cherry^+^) and that stain positive for the proliferation marker Ki67^+^ (black fraction) in comparison with cells negative for Ki67^−^ (white fraction). (D-H) MCF7 breast cancer cells expressing a Cre recombinase construct (Cre^+^ MCF7) were treated with DMSO or 500nM Palbociclib (Palbo) for 10 days to induce senescence. Exosomes were purified from Cre^+^ MCF7 cells (Cre-ex) and used to treat MCF7 cells expressing a fluorescent reporter gene which switches from expressing dsRED to eGFP upon exosome internalization (Reporter-MCF7). (E) Representative pictures showing exosome uptake (green cells) in Reporter MCF7s treated with exosomes isolated from Cre^+^ MCF7 treated with DMSO or Palbo. Staining with Sudan Black show that the GFP^+^ cells are also positive for Sudan Black staining. (F) Quantification of the percentage of GFP^+^ Reporter MCF7s treated with exosomes presenting Sudan Black staining. (G) Pictures showing exosome uptake (GFP^+^) in MCF7 reporter cells incubated with Cre-ex from DMSO or Palbo treated cells. Arrow shows that GFP^+^ cells treated with Cre-ex from DMSO cells are also positive for Ki67, while GFP^+^ cells incubated with Cre-ex from Palbo treated cells are negative for Ki67. (H) Quantification of the percentage of GFP^+^/Ki67^+^ Reporter-MCF7 treated with exosomes purified from DMSO or Palbo-treated Cre+MCF7. (E,G) Scale bar, 100μm. **See also Figure S3.**

### Cre-loxP reporter system confirms a direct link between exosome uptake and induction of senescence

To further confirm a direct role for exosomes derived from senescent cells in inducing paracrine senescence, we next used the Cre *loxP* reporter system previously described (Zomer et al., 2015; Zomer et al., 2016). For this, we generated MCF7 breast cancer cells expressing Cre recombinase (Cre^+^ MCF7) and MCF7s expressing a *loxP* flanked DsRed construct (Reporter MCF7) (**Figure S3B**). As previously shown with HFFF2s, exosomes isolated from MCF7s undergoing senescence by treatment with 500nM of the CDK4/6 inhibitor, Palbociclib (Palbo), for 10 days (Rapisarda et al., 2017) induced senescence in normal MCF7 cells, as shown by the decrease in cell number and an increase in SA-p-Gal staining and the mRNA levels of *IL-8* and *TP53* (**Figure S3C-E**). The incubation of Reporter MCF7 with exosomes expressing Cre (Cre-ex) from both DMSO and Palbo-treated MCF Cre^+^ cells, switch from expressing DsRed to enhanced GFP (Figure 3D,E). Furthermore, by measuring the percentage of GFP^+^ MCF7 Reporter cells (that have taken up Cre-ex) we could observe a positive correlation between GFP^+^ and Sudan Black^+^ cells, which stains for lysosomal aggregates, a characteristic of senescence (Georgakopoulou et al., 2013) (Figure 3E). In fact, in Figure 3F we show that the percentage of GFP^+^/Sudan Black^+^ cells is higher upon treatment with Cre-ex isolated from Palbo-treated MCF7s. Additionally, we could observe that GFP^+^ cells were negative for Ki67 when treated with Cre-ex derived from Palbo treated cells (Figure 3G) and that the percentage of GFP^+^/Ki67^+^ cells decreased upon treatment with Palbo- derived Cre-ex (Figure 3H). Altogether, these data confirm that exosomes derived from senescent cells are indeed the direct cause of paracrine senescence induction.

### Human and *ex vivo* mouse cells undergoing senescence secrete more exosomes

Previous studies have shown that cells undergoing senescence release more exosomes (Kavanagh et al., 2017; Lehmann et al., 2008; Takasugi et al., 2017). Thus, we quantified the number and size of exosomes by nanoparticle tracking analysis (NTA) in HFFF2 undergoing OIS and DDIS and show an increase in exosome release during senescence (Figure 4A). Furthermore, breast primary fibroblasts expressing ER:RAS, MCF7 cells treated with 500nM Palbo and mouse Hepatic Stellate Cells (mHSC) derived from an adult mouse harboring a doxycycline (Dox) inducible construct to express shp53 (Krizhanovsky et al., 2008; Lujambio et al., 2013) also showed an increase in the secretion of exosomes upon the induction of senescence (**Figure S4A-C**). Next, we characterized the exosome population released in iRAS cells by capturing exosomes onto beads coated with a CD63 antibody, followed by a CD81-PE conjugated antibody as in **Figure S1A**. In fact, at different time points after inducing senescence we can see an increase in CD81 fluorescence intensity and therefore Cd63^+^/CD81^+^ exosome release (Figure 4B). Altogether, our results show that the activation of the senescence program induces exosome release in mouse and human cells.

**Figure 4.**
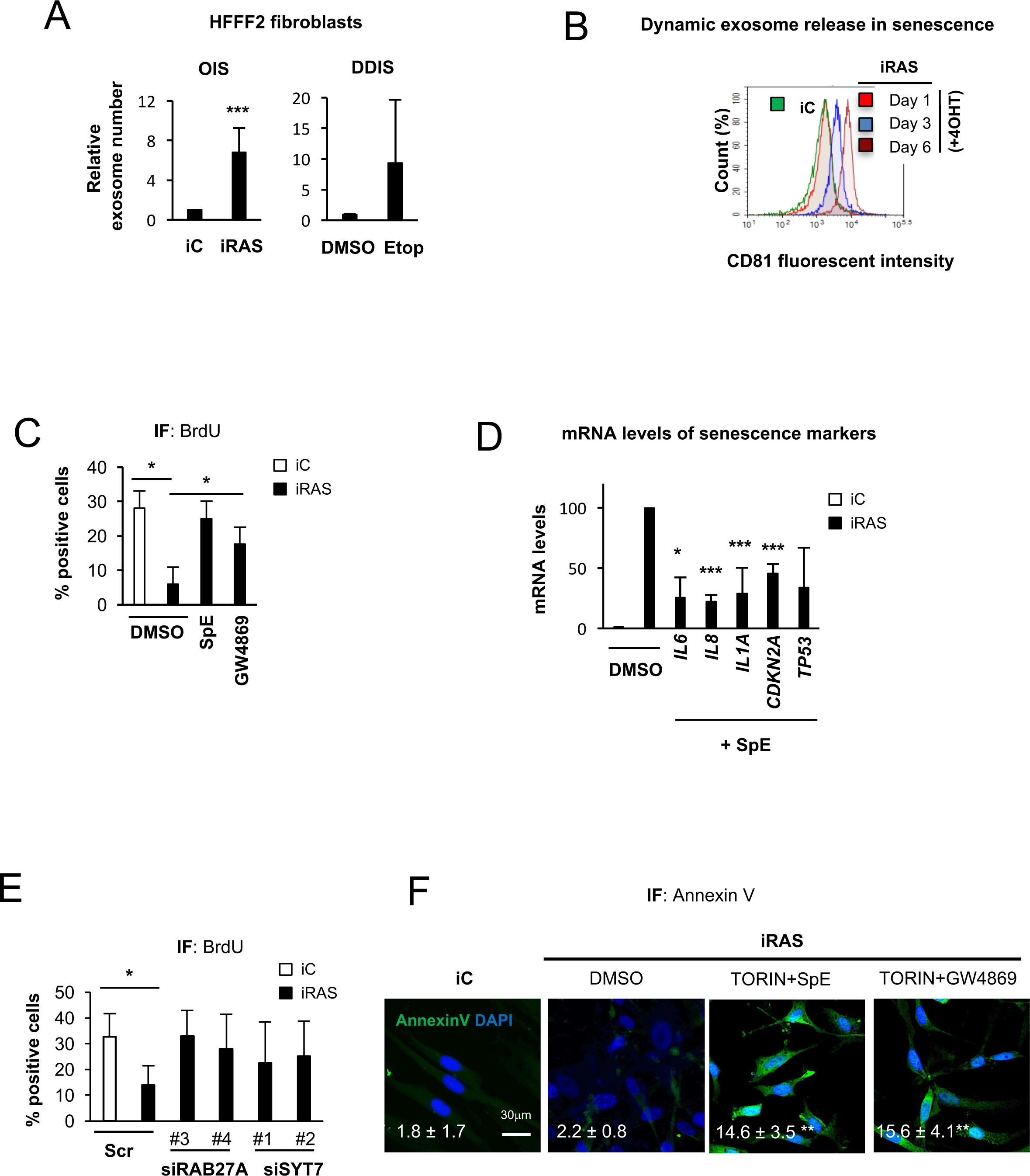
Exosome biogenesis is an essential biological process to induce senescence. (A) The number of exosomes released by HFFF2 undergoing OIS and DDIS was determined using nanoparticle tracking analysis (NTA). OIS and DDIS was established by treating iRAS with 200nM 4OHT (OIS) or HFFF2 fibroblasts with 50μM Etoposide (DDIS) for 2 days followed by 5 days with fresh 0.5% FBS-medium. Data represents the mean ± SD of 2-5 independent experiments. (B) The number of CD63+/CD81 + vesicles at different time points (1, 3, 6 days) after the induction of senescence (+4OHT) in iRAS HFFF2s was determined by FACS. (C,D) iRAS HFFF2s were treated with 10 μM GW4869 or 5 μM SpE for 72h and (C) IF was performed by staining for BrdU to determine proliferation or (D) RNA was isolated to determine the mRNA expression levels of *IL6, IL8, IL1A, CDKN2A* or *TP53* in iRAS HFFF2 treated with SpE. Data are normalized to iRAS cells treated with 4OHT and represent the mean ± SD of (C) 4 and (D) 2-4 independent experiments. (E) iRAS HFFF2s were transfected with either a Scr or siRAB27A or siSYT7 for 3-5 days. Subsequently, cells were fixed and stained to determine the proliferation rate by measuring BrdU incorporation. (F) IF for Annexin V in iRAS HFFF2s treated simultaneously with Torin-2+GW4869 and Torin-2+SpE shows an increase in nuclear staining upon treatment with both inhibitors. Data are showing the percentage of cells with nuclear staining from 2 independent experiments. Scale bar, 30μm. **See also Figure S4.**

### Exosome biogenesis is an essential feature of senescent cells

Next, we wanted to ask if exosome biogenesis is an essential process during senescence. To test this, we treated iRAS HFFF2 with SpE and GW4869 and determined whether blocking exosome biogenesis affected senescence. To our surprise and in contrast with published data (Kavanagh et al., 2017; Takahashi et al., 2017), blocking exosome biogenesis prevented the establishment of senescence as shown by a partial reversion in the percentage of HFFF2s incorporating BrdU (Figure 4C) and a reduction in the levels of p-γH2AX positive HFFF2 by IF (**Figure S4D,E**). Furthermore, qPCR analysis to determine the mRNA levels of additional markers of senescence also show a partial reduction in the upregulation of *IL6, IL8, IL1A, CDKN2A* and *CDKN1A* when iRAS cells were treated with either SpE or GW4869 (Figure 4D, **Figure S4F**). We also observed that the use of two independent siRAB27A and siSYT7 prevented the cell cycle arrest characteristic of senescence (Figure 4E). In fact, iRAS HFFF2 expressing a vector encoding for two previously characterized short hairpin RNA targeting p53 (shp53) and p16^INK4A^ (shp16) (Acosta et al., 2008; Rapisarda et al., 2017) show a reduction in the release of exosomes when RAS is expressed (+ 4OHT) (**Figure S4G**). Importantly, the simultaneous use of inhibitors of the SASP (Torin-2) and inhibitors of exosome biogenesis (SpE or GW4869) induced cell death in HFFFs as shown by nuclear staining of the apoptosis marker Annexin V (Figure 4F) suggesting that both pathways are equally essential for the successful establishment of the senescence program in primary fibroblasts. These data show that blocking exosome biogenesis partially prevents the induction of senescence in HFFF2 cells.

### Exosome protein cargo is responsible for inducing paracrine senescence

Based on our previous finding, we hypothesized that the protein content of the exosomes derived from control and senescent HFFF2s might differ. For this, we subjected purified exosomes from iRAS and Etop-treated HFFF2 and their respective controls to label-free mass spectrometry (MS) analysis (Figure 5A). GO analysis of the 1600 proteins identified show the Cellular component “Extracellular exosome” is overrepresented (Figure 5B). To identify proteins exclusively regulating senescence, we selected proteins deregulated in both OIS- and DDIS-exosomes *vs* control exosomes with > 2 log_2_ fold change difference and adjusted false discovery rate (FDR) < 0.01 in both OIS and DDIS. The Venn diagram analysis shows that 265 proteins are deregulated during both OIS and DDIS (Figure 5C), which group into GO Biological Processes related to senescence, as “Wound healing” and “Cell adhesion” (Figure 5D). Furthermore, volcano plot analysis comparing individually OIS- and DDIS-exosomes protein *vs* their respective controls show that most proteins identified within exosomes derived from senescent HFFF2s are upregulated (**Figure S5A**). To determine which of these proteins are essential to mediate paracrine senescence, we conducted a small-scale screen using siRNA SMART pool in iRAS cells for the top 50 proteins most upregulated in both DDIS and OIS (>2 log_2_ fold change; p<0.05; > 4 peptides fold difference between control and senescence) (**Figure S5B**). We used the CM from iRAS to determine paracrine senescence in normal HFFF2 (Figure 5E) using siRAB27A and siSYT7 as positive controls (green bars) and determined BrdU incorporation and p21^CIP^ protein levels by IF (Figure 5F; **Figure S5C**). The most consistent candidates preventing both growth arrest and upregulation of p21^CIP^, in addition to an increase in p16^INK4A^, IL-8 and p-γH2AX by IF (**Figure S5D**), are two proteins implicated in interferon (IFN) signaling: interferon-inducible transmembrane protein 3 (IFITM3) and interferon-induced GTP-binding protein (MX1) (red bars). Here, we show that the protein content from control or senescent derived exosomes is different and identify IFITM3 and MX1 as essential in inducing paracrine senescence.

**Figure 5.**
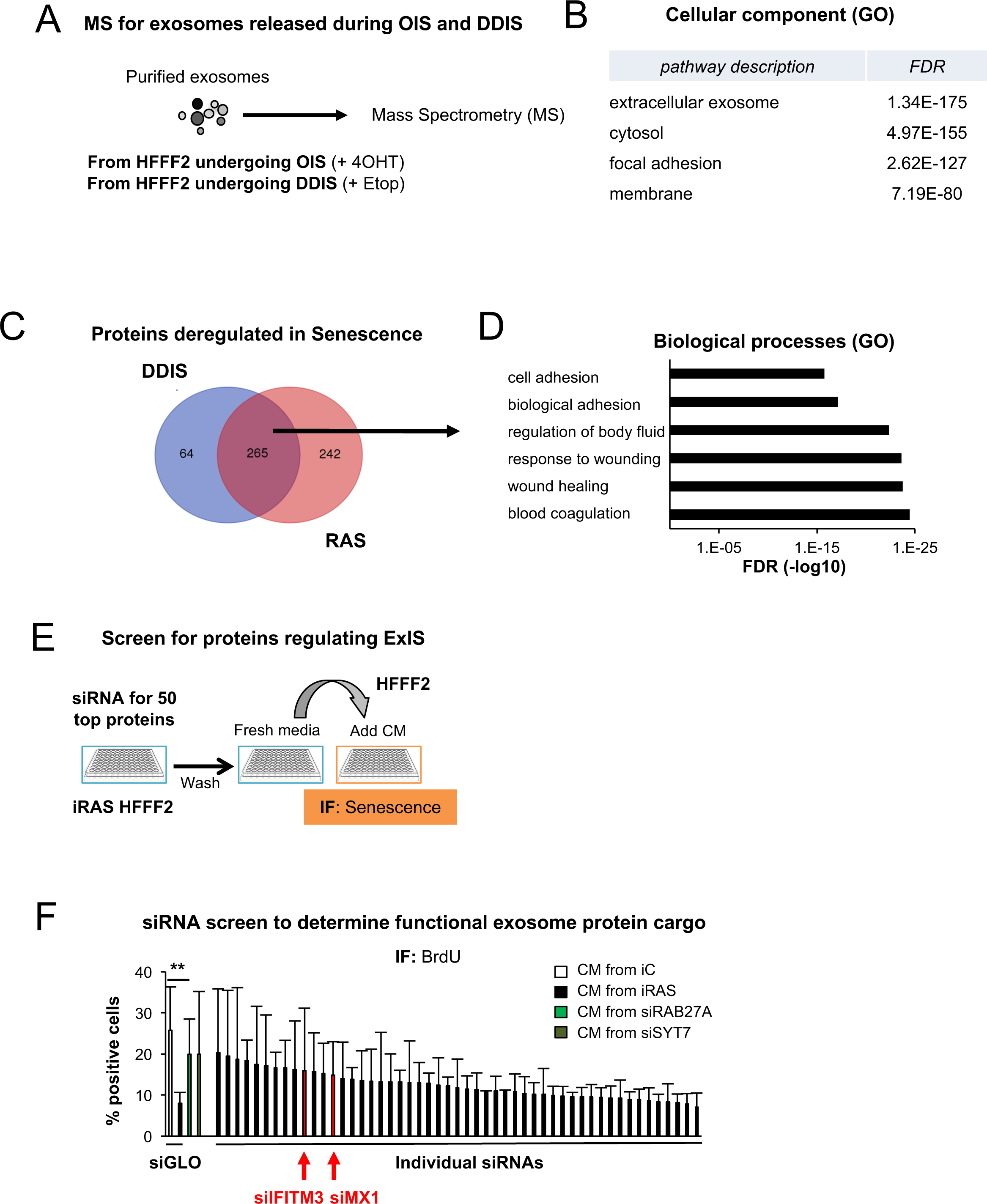
MS proteomic analysis reveal a specific cargo in exosomes derived from senescent cells. (A) Scheme showing the Mass Spectrometry (MS) approach. Exosomes were isolated from HFFF2 undergoing either OIS or DDIS from two independent experiments and were sent for label free MS analysis. (B) DAVID GO analysis for the 1600 proteins detected by MS group into the “Extracellular exosome” pathway. FDR (false discovery rate). (C) Venn diagram for proteins with >2 log2 differential expression and < 0.01 FDR in exosomes released during OIS and DDIS compared to control show 265 proteins in common being deregulated during senescence. (D) GO analysis using STRING bioinformatics groups the 265 proteins differentially regulated in both OIS and DDIS into Biological Processes related to senescence like “Wound healing” and “Response to Wound Healing”. (E) Schematic diagram showing the strategy used for the performance of the siRNA screen. Briefly, after transfection with the siRNA the CM was washed out and replenished with fresh media for 72h as in Figure 1. (F) Screen using SMART pool siRNA targeting the 50 most upregulated proteins in both OIS and DDIS with > 4 peptide fold difference between control and senescence. Data show BrdU staining by IF for HFFF2s treated with different CM. siRAB27A and siSYT7 were used as positive controls (green bars). siIFITM3 and siMX1 are highlighted in red. Data represent the mean ± SD of 3 independent experiments. **See also Figure S5.**

### IFITM3 is essential for paracrine senescence mediated by exosomes

Next, we decided to investigate the influence of IFITM3 within exosomes on mediating paracrine senescence. For this, we depleted IFITM3 from iRAS cells and isolated exosomes from these cells. Immunoblotting analysis of the levels of IFITM3 in exosomes shows an increase in IFITM3 within exosomes derived from iRAS cells which is lost when cells are transfected with siIFITM3 (Figure 6A). However, no changes in the total protein levels of either IFITM3 nor MX1 could be observed in the donor cells lysates from iRAS cells (Figure 6B, **Figure S6A**). Importantly, treatment of HFFF2 with an equal number of exosomes derived from iC and iRAS transfected with Scr or siIFITM3 shows that the upregulation of p16^INK4A^ protein levels by IF is dependent on the presence of IFITM3 in exosomes (Figure 6C). Interestingly, NTA analysis of iC and iRAS cells transfected with and without siIFITM3 show no differences in exosome release (**Figure S6B**) confirming that the presence of IFITM3 in the exosomes, and not exosome number, is responsible for mediating paracrine senescence. Furthermore, analysis of the RNA transcripts deregulated upon incubation of HFFF2 with exosomes derived from OIS- and DDIS-cells (Figure 2E) show that the IFN pathway is deregulated upon exosome treatment (Figure 6D,E, **Figure S6C**), with several *IFITM*, the related IFN-induced protein with tetratricopeptide repeats *(IFIT)* and *MX1-2* transcripts being deregulated in HFFF2 treated with senescent exosomes but not when exosome biogenesis is prevented with SpE (Figure 6D,E). However, no changes where observed in *IFN* or *IFN receptors (IFNR)* transcripts that where common to both OIS and DDIS, as previously described (Dou et al., 2017). Additionally, we could not detect secretion of IFNg to the CM from either iC or iRAS cells (Figure 6F) or detect differences in protein levels for IFN or IFNR in exosomes in the MS data. Yet, we could detect IL-8 secretion in iRAS cells (Figure 6F). Finally, simultaneous treatment of HFFF2 with exosomes derived from iRAS and the classical mediator of IFN signaling, JAK1/2 inhibitor 1μ;M (Ruxolitinib) does not prevent the activation of senescence, while an inhibitor specific for p38MAPK (SB202190) prevented the growth arrest induced by senescent exosomes. Altogether, these data suggest that IFITM3 is essential for mediating paracrine senescence possibly by activating a non-canonical IFN pathway in recipient cells.

**Figure 6.**
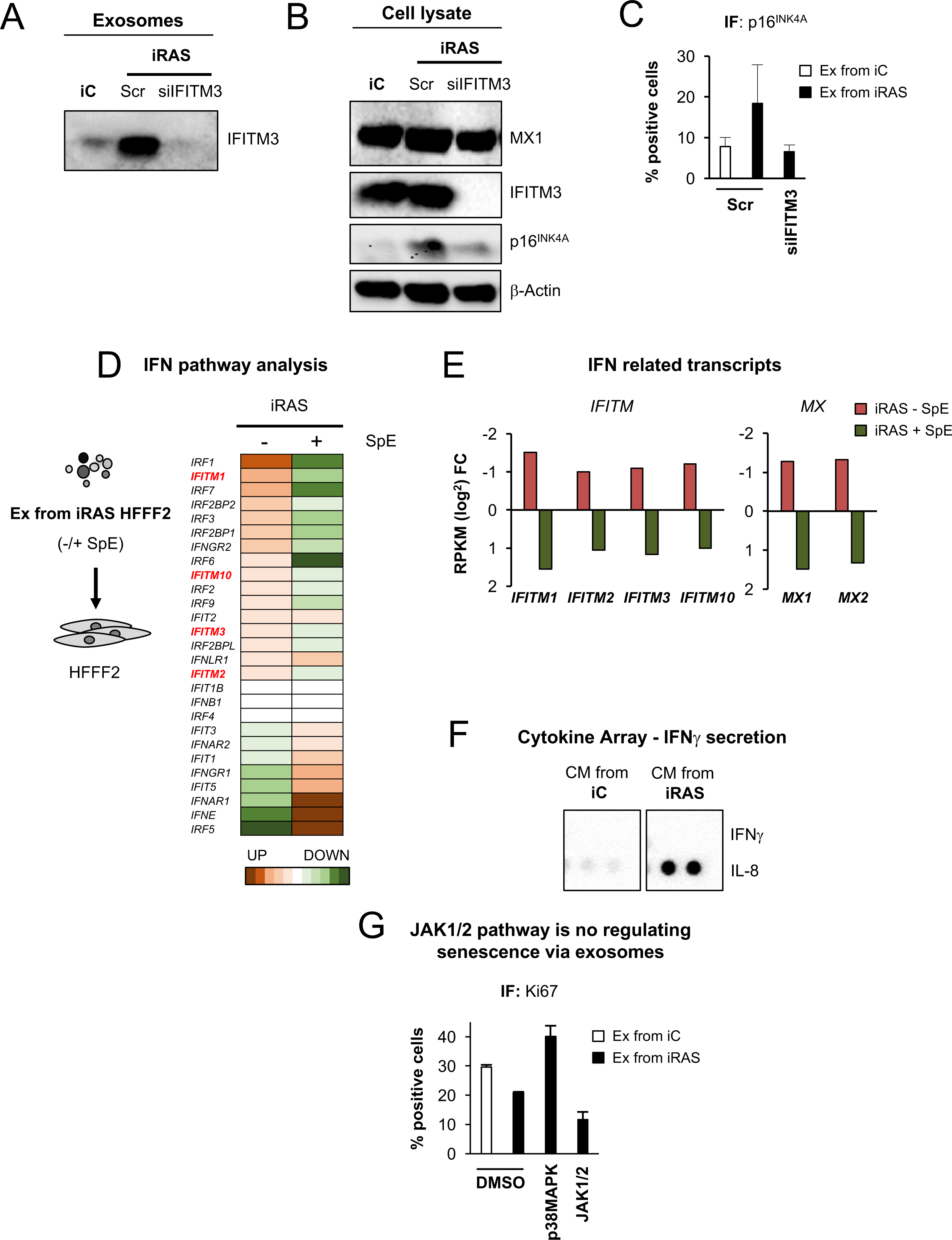
IFITM3 within exosomes is essential to induce paracrine senescence. (A) Immunoblotting for IFITM3 present in exosomes derived from iRAS transfected with or without an siRNA targeting IFITM3 (siIFITM3). An equal number of exosomes was loaded. (B) Immunoblotting analysis for cell lysates derived from iRAS HFFF2 transfected with Src or siIFITM3. β-Actin is the loading control. (C) IF staining for p16^INK4A^ in HFFF2s treated with the same number of exosomes derived from iC or iRAS transfected with or without siIFITM3. (D,E) HFFF2s treated with exosomes derived from iRAS cells shows an increase in transcripts related to the interferon (IFN) pathway, in particular *IFITM* (in red) and *IFIT* mRNAs, which are downregulated when HFFF2s are treated with exosomes isolated from iRAS HFFF2 treated with SpE. (E) *IFITM* and *MX* transcripts are specifically shown. Data has been normalized to the control and represents the mean ± SD of 3 independent experiments (RPKM log_2_ fold difference). (F) The secretion of IFNg was determined by semi-quantitative cytokine array dot blot in iC and iRAS CM. IL-8 secretion was observed in the CM of iRAS and used as positive control that the array worked. A representative blot of 2 independent experiments is shown. (G) HFFF2 were treated simultaneously with exosomes isolated from iRAS cell and DMSO, 1μM JAK1/2 (Ruxonitilib) or 20μM p38MAPK inhibitor (SB202190) for 72h. Ki67 was used to determine proliferation. **See also Figure S6.**

## DISCUSSION

Intercellular communication is an important mechanism by which cells interact with each other. It can be mediated in the form of soluble factors or extracellular vesicles (Kuilman and Peeper, 2009; Tkach and Thery, 2016). However, in the context of senescence, a role for extracellular vesicles has been neglected, although some studies have found the “Extracellular Vesicle Pathway” over-represented by GO analysis of differentially expressed genes in senescence (Hoare et al., 2016; Lujambio et al., 2013). Furthermore, recent studies have highlighted the importance of exosome biogenesis for the cellular homeostasis (Takahashi et al., 2017) and for promoting cancer cell proliferation (Takasugi et al., 2017).

Non-cell autonomous (paracrine) senescence via the SASP has been previously described as an important mechanism for the clearance of senescent cells by the immune system (Acosta et al., 2013; Dou et al., 2017; Hoare et al., 2016), although these studies do not discern between the effect of soluble factors and EVs. Here, we are providing evidence that both the soluble fraction and exosomes are responsible in mediating paracrine senescence. We show not only that blocking exosome biogenesis using small molecule inhibitors and RNAi prevent the induction of senescence but that treatment of normal fibroblasts and cancer cells with exosomes-derived from senescent cells using a variety of stimuli induce senescence. Furthermore, we observe a direct correlation between exosome-uptake and senescence activation in different cell lines with two independent triggers of senescence. Altogether, we demonstrate that exosomes are responsible for mediating paracrine senescence and speculate they play an important role in inducing bystander senescence in therapy-induced senescence (Demaria et al., 2017), or in mediating bystander aging (Acosta et al., 2013; Nelson et al., 2012). In fact, when compared to soluble factors, exosomes have very different biophysical and biochemical properties. Exosomes have a longer-lifespan than soluble factors as they are more resistant to protease degradation and can mediate both local and systemic signaling (O’Loghlen, 2018). Interestingly, in the luciferase knock-in mouse model *(p16*^*LUC*^) activation of *p16Ink4a*^*+*^ cells was observed not only in emerging neoplasias, but also in the nearby stroma, suggesting that senescence can be transmitted via non-autonomous mechanisms (Burd et al., 2013). This leads us to speculate that indeed paracrine senescence mediated by exosomes can be transmitted *in vivo*, although more functional experiments would be needed to confirm this.

Exosome biogenesis is an important feature of many cells and a recent study has shown its relevance in maintaining cellular homeostasis (Takahashi et al., 2017). In contrast with our data, the authors show a DNA-damage response upon inhibition of exosome biogenesis, which we only observe when blocking both the release of soluble factors and exosomes, which could be due to different experimental settings. However, a common finding is that fact that most cells undergoing senescence triggered by a variety of stimuli release more exosomes in human and mouse cells (Kavanagh et al., 2017; Lehmann et al., 2008; Takahashi et al., 2017). In addition, we observed that blocking exosome release with small molecule inhibitors and RNAi prevents the activation of senescence (although to a lesser extent than paracrine signaling), highlighting the importance of this pathway during senescence.

Recently, many studies have found a cellular response characteristic of infectious agents during senescence in the absence of pathogens. For example, it has been show that the SASP is regulated by the inflammasome via IL-1a signaling (Acosta et al., 2013) and by the cGAS-STING (cyclic GMP-AMP synthase linked to stimulator of interferon genes) pathway (Dou et al., 2017; Gluck et al., 2017; Yang et al., 2017). Both pathways are induced during senescence in the absence of pathogens or in the presence of doublestranded DNA. Here, we show that the non-canonical IFN pathway (Diamond and Farzan, 2013; Platanias, 2005) is being activated by exosomes derived from senescent cells in the absence of pathogens, as we don’t observe differences in IFNγ secretion between iC or iRAS nor detect IFN or IFNR within exosomes by MS analysis. In fact, HFFF2 treated with exosomes derived from iRAS upregulate *IFITM* and *IFIT* transcripts without significant changes in *IFN* and *IFNR* RNA common to both OIS and DDIS. Interestingly, IFITM3 is found enriched in exosomes derived from iRAS HFFF2s without being affected in the donor cells undergoing senescence. Finally, blocking the JAK1/2 pathway via small molecule inhibitors, which is activated upon IFN signaling, does not prevent senescence mediated by exosomes. This is not an unprecedent finding, as exosomal RNA from stromal cells has been shown to stimulate the viral RNA pattern recognition receptor (PRR) RIG-I and IFN pathway in the absence of viral RNA (Boelens et al., 2014; Nabet et al., 2017). However, at present we cannot exclude that DNA/RNA present as cargo or attached to the exosomes are responsible for stimulating IFN signaling, as other reports have found IFN-mediated signaling via exosomes (Cossetti et al., 2014; Li et al., 2013), a role for *IFNAR1* during OIS (Katlinskaya et al., 2016) and DNA from senescent cells secreted via exosomes (Takahashi et al., 2017).

Although our data exclude a role for microvesicles during senescence, and here we are describing exosomes as a heterogeneous population, it would be interesting for future studies to address whether small extracellular vesicles of endocytic origin or not mediate a similar cellular response in normal cells depending on specific populations (Tkach et al., 2018).

Finally, the idea that blocking exosome secretion could be a potential therapeutic approach to alleviate senescence “spreading” during senescence-induced by chemotherapy treatment or accumulation in aging tissues presents itself as a very attractive tool in the future.

## AUTHORS CONTRIBUTIONS

MB performed most of the experiments, with help from JFL and PCF. HP provided advise with the proteomic experimental settings. PXE and JM performed the proteomics analyses. AO conceived and designed the study. AO wrote and edited the manuscript, with input from all the authors.

## ACKNOWLEDGEMENTS

We are grateful to Tom Nightingale, Maria Niklison-Chirou and lab members for critical reading of the manuscript; Alissa Weaver for providing fluorescently tagged-CD63 lentiviral constructs; Luke Gammon for excellent technical advice with the In Cell; Belen Martin for assistance with the Confocal and Airyscan microscopy; Rob Lowe and QMUL Genome Centre for the RNA sequencing and data analysis; Gary Warnes for assistance with the FACS. Mouse hepatic stellate cells were a kind gift from Scott Lowe.

AO’s lab is supported by the BBSRC (BB/P000223/1) and MB is funded by the MRC (MR/K501372/1). PCF and JFL are funded by the Xunta de Galicia Fellowships (Spain).

## STAR METHODS

## CONTACT FOR REAGENT AND RESOURCE SHARING

“Further information and requests for resources and reagents should be directed to and will be fulfilled by the lead contact, Ana O’Loghlen.”

## EXPERIMENTAL MODELS

### Cell culture

HFFF2 human foreskin primary fibroblasts were obtained from the Culture Collections (Public Health England, UK). IMR-90, MCF7 and HEK293T were bought from ATCC. Breast fibroblasts (BF) were isolated from a breast mastectomy and have been described elsewhere (Rapisarda et al., 2017). All cells were grown in high glucose Dulbecco’s modified Eagle’s medium with 10% fetal bovine serum and 1% antibiotic-antimycotic solution. Mouse hepatic stellate cells were a kind gift from Scott Lowe and were grown in 1μg/ml of Doxycycline.

## METHOD DETAILS

### Soluble fraction, exosome and microvesicles isolation

All cells were maintained in exosome-depleted FBS. FBS was depleted of exosomes by overnight ultracentrifugation at 100,000 g at 4°C (Sorvall 100SE Ultracentrifuge). The supernatant was removed and stored in 50 mL falcons at −20°C until required. All CM was collected after 72 h incubation with cells. Whole CM was centrifuged at low speed to eliminate dead cell and cellular debris prior to use.

To isolate exosomes, the protocol of differential ultracentrifugation (Thery et al., 2006) was modified and adapted. The CM was filtered through a 0.22 μm filter prior to the 100,000g centrifugation step and also after in some cases (for the Cre-LoxP experiments). MV were collected after the 10,000g centrifugation step. The supernatant of the CM was collected after the 100,000g centrifugation step and concentrated using a 10K column (Amicon™ Ultra-0.5 Filter). The final 100,000 g pellet was washed twice in 15 ml of PBS and resuspended in 100 μl of sterile DMEM for the functional cell culture experiments. Alternatively, for Western Blot analysis, exosome pellet was re-suspended in protein lysis buffer. A Sorvall 100SE Ultra Centrifuge, with a Beckmann Fixed Angle T865 rotor was used for all exosome isolations.

### CM with siRNA and inhibitors functional experiments

24h after plating HFFF2 in 96-well plates, senescence was induced by adding 200nM 4OHT. The following day the cells were washed and supplemented with 0.5% FBS and the indicated inhibitors for 2-3 days, after which plates were washed again to remove the inhibitors from the media and supplemented with fresh media (0.5% FBS) for further 72h. For the experiments with the siRNA, reverse transfection with 30nM siRNA was performed and the media replenished (10% FBS) with 4OHT after 2 days and left for an additional 48h. After, the cells were washed to remove all siRNA from the media and incubated with 0.5% FBS fresh media for an additional 72h. For the mini-screen 50nM of SMARTPool siRNA was used.

### Treatment of cells with CM or isolated exosomes

Fibroblasts were treated with the CM from several experiments previously centrifuged at low speed to discard dead cells and supplemented with FBS to reach 10% in the final volume. HFFF2 treated with isolated exosomes were also supplemented with media containing 10% FBS, while in the experiment performed with MCF the cells were incubated with isolated exosomes resuspended in 0.5% FBS media.

### Nanoparticle tracking analysis (NTA)

Prior to the NTA analysis, the NanoSight LM10 equipped with a 405nm laser (Malvern Instruments) was calibrated using Silica Microspheres beads (Polyscience). Samples to be measured were then diluted in PBS in order to obtain a particle number between 10^8^-10^9^ particles. At least three repeated measurements of 60s were taken per each individual sample and the mean value was used to determine particle number. Static mode (without flow) was used for each analysis. The movement of each particle in the field of view was measured to generate the average displacement of each particle per unit time which was calculated using the NTA 3.0 software.

### Affinity-based capture of exosomes on beads

For exosome characterization by flow cytometry, aldehyde/sulphate latex beads (Thermo Fisher) were coated with anti-CD63 antibody and incubated with the different CM overnight at 4C in a rotations wheel. After extensive washing, anti-CD81-PE conjugated antibody was added for further 40min at RT, washed with PBS and acquired using NovoCyte Flow Cytometer (Acea, Biosciences).

### β-Galactosidase staining

Cells were washed with PBS and fixed with 0.05% (w/v) glutaraldehyde (in PBS) for 15 mins at RT. Cells were washed a second time with PBS and incubated with 5-bromo-4- chloro-3-indolyl-beta-D-galacto-pyranoside (X-gal) solution for 12 hr at 37C. Cells were imaged at 4 and 6 hr using a light microscope (Nikon) at 20X magnification and single representative images of each well were taken.

### Immunofluorescence staining

Cells grown in 96-well plates were washed with PBS and fixed in 4% paraformaldehyde for 15 minutes at RT. Cell were then washed in PBS twice before been permeabilized and blocked for 40 minutes with 0.2% Tritonx100 together with 1% BSA and 0.2% gelatin fish (Sigma). For IF staining, cells were incubated overnight with the primary antibody and in the case of BrdU detection treated with DNaseI and MgCl_2_. Cells were then washed in PBS and then incubated one hour with secondary antibody, DAPI and Cell Mask Deep Red (Invitrogen). Immunofluorescence images were acquired using IN Cell 2200 automated microscope (GE)

### RNA extraction, cDNA synthesis and qPCR

Cells grown in 6-well plates or 10 cm dishes were washed with PBS and lysed directly into the culture dish using TRIzol Reagent (Thermo Fisher). cDNA synthesis was performed using the High-Capacity cDNA Reverse Transcription Kit (Thermo Fisher). qPCR reactions were performed using SYBR Green PCR Master Mix (Applied Biosystems,) on a 7500 Fast System RealTime PCR cycler (Applied Biosystems). Primer sequences are listed in **Table S1**.

### Gene expression

Stable retroviral and lentiviral expression was performed as in previous studies (Acosta et al., 2008; Rapisarda et al., 2017).

### Protein analysis by immunoblotting

Exosomes were lysed using the following buffer lysis buffer [(Tris-HCl 20 mM pH 7.6; DTT 1 mM; EDTA 1 mM; PMSF 1 mM; benzamidine 1 mM; sodium molybdate 2 mM; β- sodium glycerophosphate 2 mM; sodium orthovanadate 0.2 mM; KCl 120 mM; 1 μg/ml (each) de leupeptine, pepstatine A and antipaína; NonidetTM P-40 0.5% (v/v); Tritón X- 100 0,1% (v/v)], separated in an SDS-PAGE gel, transferred to a PVDF membrane and probed with different antibodies.

### RNA sequencing

For RNA-seq, total RNA was extracted using TRIzol Reagent. First strand cDNA synthesis was performed with superScript III First-Strand Synthesis System. After purification using SPRI beads, the double stranded cDNA was ligated to in house designed adapters (based on TruSeq Indexed adapters (Illumina)) using NEBNext Ultra II (NEB) followed by 15 cycles of amplification and library purification. Sequencing was performed on an Illumina NextSeq500, High Output run with 75bp paired-end at the Genomics Centre (QMUL).

### MS proteomics

LC-MS/MS was done by coupling a nanoLC-Ultra 1D+ system (Eksigent) to an Impact mass spectrometer (Bruker) via a Captivespray source (Bruker) supplemented with a nanoBooster operated at 0.2 bar/min with isopropanol as dopant. Peptides were loaded into a trap column (NS-MP-10 BioSphere C18 5 μm, 20 mm length, NanoSeparations) for 10 min at a flow rate of 2.5 μl/min in 0.1% FA. Then peptides were transferred to an analytical column (ReproSil Pur C18-AQ 2.4 μm, 500 mm length and 0.075 mm ID, Dr. Maisch) and separated using a 100 min effective curved gradient (buffer A: 4% ACN, 0.1% FA; buffer B: 100% ACN, 0.1% FA) at a flow rate of 250 nL/min. The gradient used was: 0-2 min 2% B, 2-102 min 33%B, 102-112 min 98% B, 112-120 min 2% B. The peptides were electrosprayed (1.35 kV) into the mass spectrometer with a heated capillary temperature of 180 °C. The mass spectrometer was operated in a data-dependent mode (130-1600 m/z), with an automatic switch between MS and MS/MS scans using a top 20 method (threshold signal ≥ 500 counts, z ≥ 2 and m/z ≥ 350). An active exclusion of 30 sec was used. The precursor intensities were re-evaluated in the MS scan (n) regarding their values in the previous MS scan (n-1). Any m/z intensity exceeding 5 times the measured value in the preceding MS scan was reconsidered for MS/MS. Peptides were isolated using a 2 Th window and fragmented using collision induced dissociation (CID) with a collision energy of 23-56 eV as function of the m/z value.

## QUANTIFICATION AND STATISTICAL ANALYSIS

### Statistical analysis

Statistical analysis was performed using Microsoft Excel Software student’s t-test.

### qPCR gene expression

Ct values were generated using the 7500 software version 2.0.6 (Applied Biosystems). Relative gene expression was calculated using the AACt method and normalized to the housekeeping gene RPS14. The relative mRNA expression level changes were expressed as a fold change relative the control.

### IF analysis

For IF analysis, all images were analyzed using the IN Cell 2200 Developer software version 1.8 (GE) as previously (Acosta et al., 2008; Rapisarda et al., 2017).

### MS proteomics analysis

Raw files were processed with MaxQuant (v 1.5.3.30) using the standard settings against a human protein database (UniProtKB/Swiss-Prot, August 2016, 20,195 sequences) supplemented with contaminants. Label-free quantification was done with match between runs (match window of 0.7 min and alignment window of 20 min). Carbamidomethylation of cysteines was set as a fixed modification whereas oxidation of methionines and protein N-term acetylation as variable modifications. Minimal peptide length was set to 7 amino acids and a maximum of two tryptic missed-cleavages were allowed. Results were filtered at 0.01 FDR (peptide and protein level). Further statistical analysis was performed using Perseus (v1.5.5.2). A minimum of three LFQ valid values per group was required for quantification. Missing values were imputed from the observed normal distribution of intensities. Then, a two-sample Student’s T-Test with a permutation-based FDR was performed. Only proteins with a q-value<0.10 and log_2_ ratio >2 or < −2 were considered as regulated.

### RNA sequencing

Genomic mapping was performed by the Genomics Centre using Fastq files aligned to HG 19 using STAR aligner implemented in BaseSpace (RNA-Seq Alignment pipeline v 1.1.0, Illumina). BAM file outputs from STAR were annotated using Partek Genomic Suite (v6.6) and the RefSeq data base (RefSeq 21). Differential analysis was performed with Partek Genomic Suite (v6.6) running ANOVA.

## DATA SOFTWARE AND AVAILABILITY

### Gene Ontology Analysis

Gene Ontology Analysis was performed using the STRING: functional protein association networks (https://string-db.org/) for the proteomic dataset. For the RNA seq datasets a combination of DAVID Functional Annotation Bioinformatics Microarray Analysis (https://david.ncifcrf.gov/) and PANTHER - Gene List Analysis (http://www.pantherdb.org/).

---

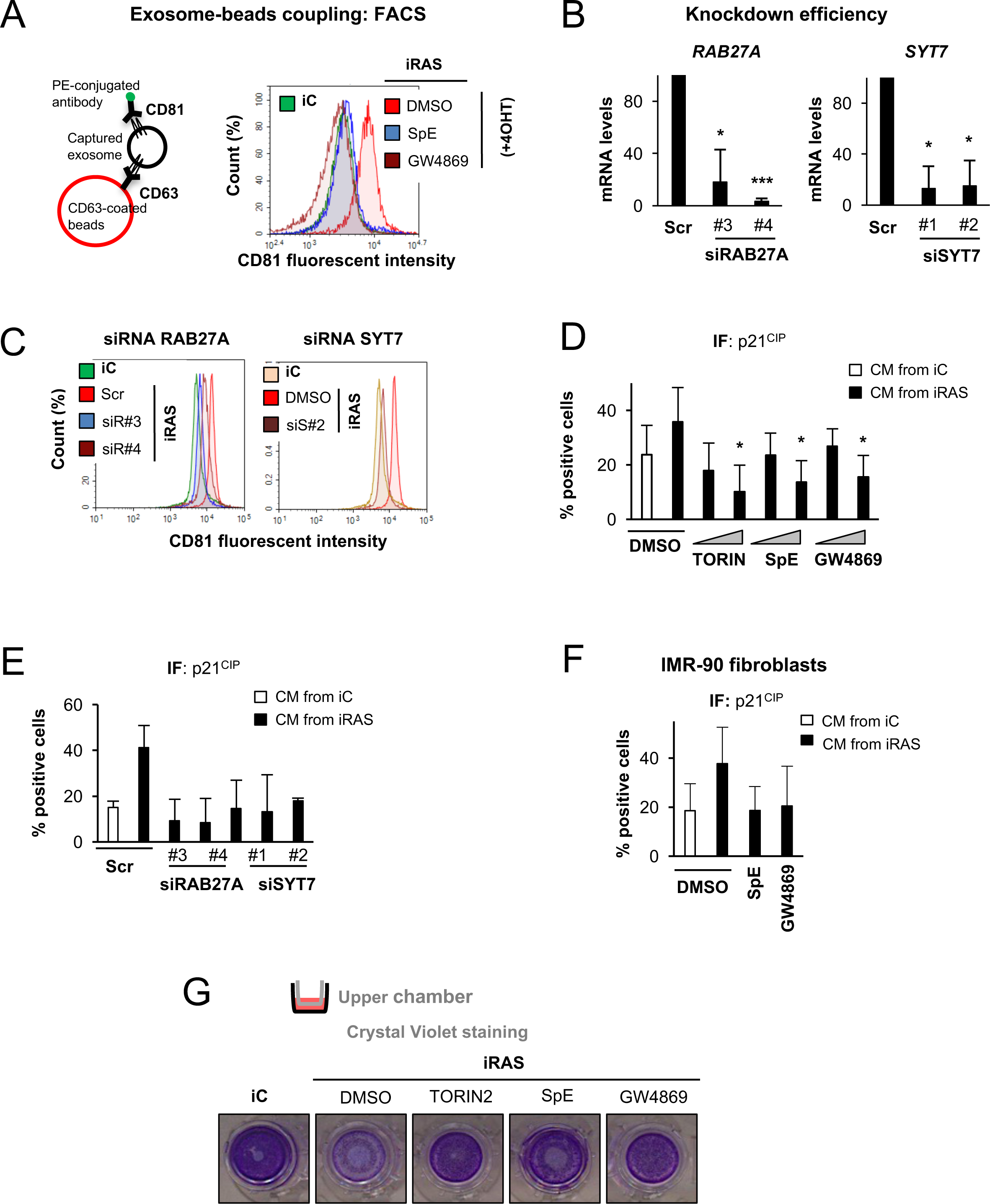

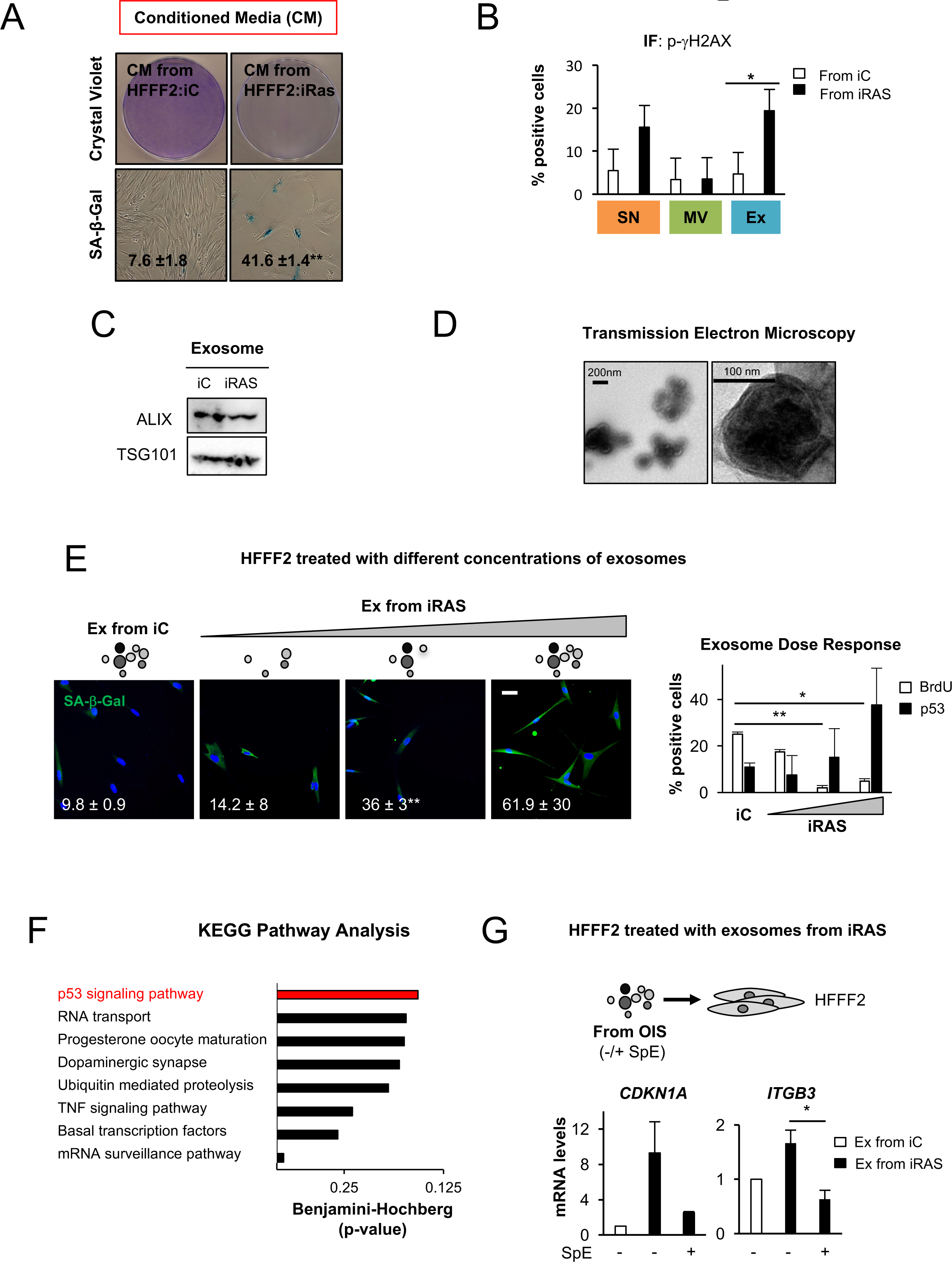

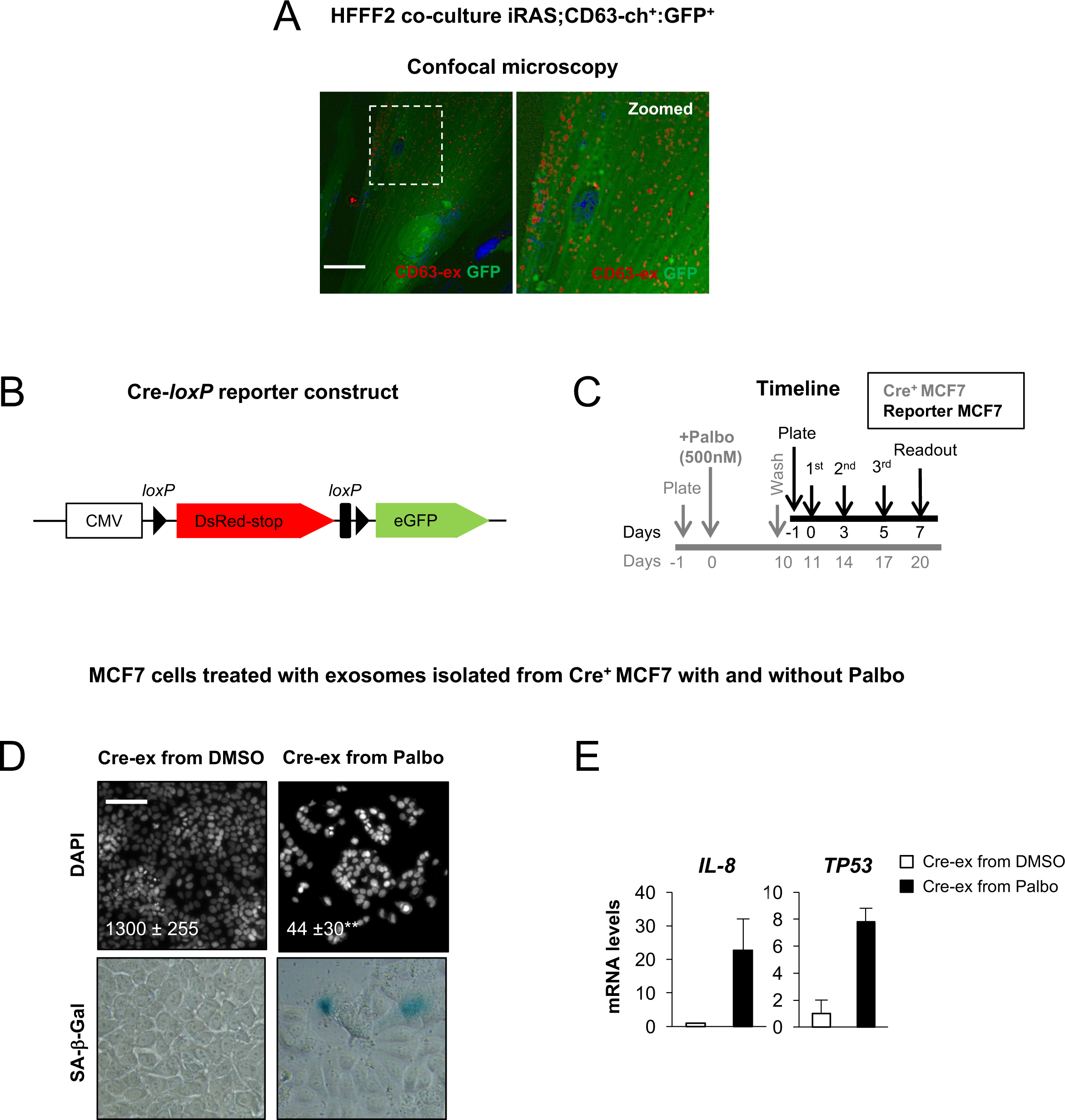

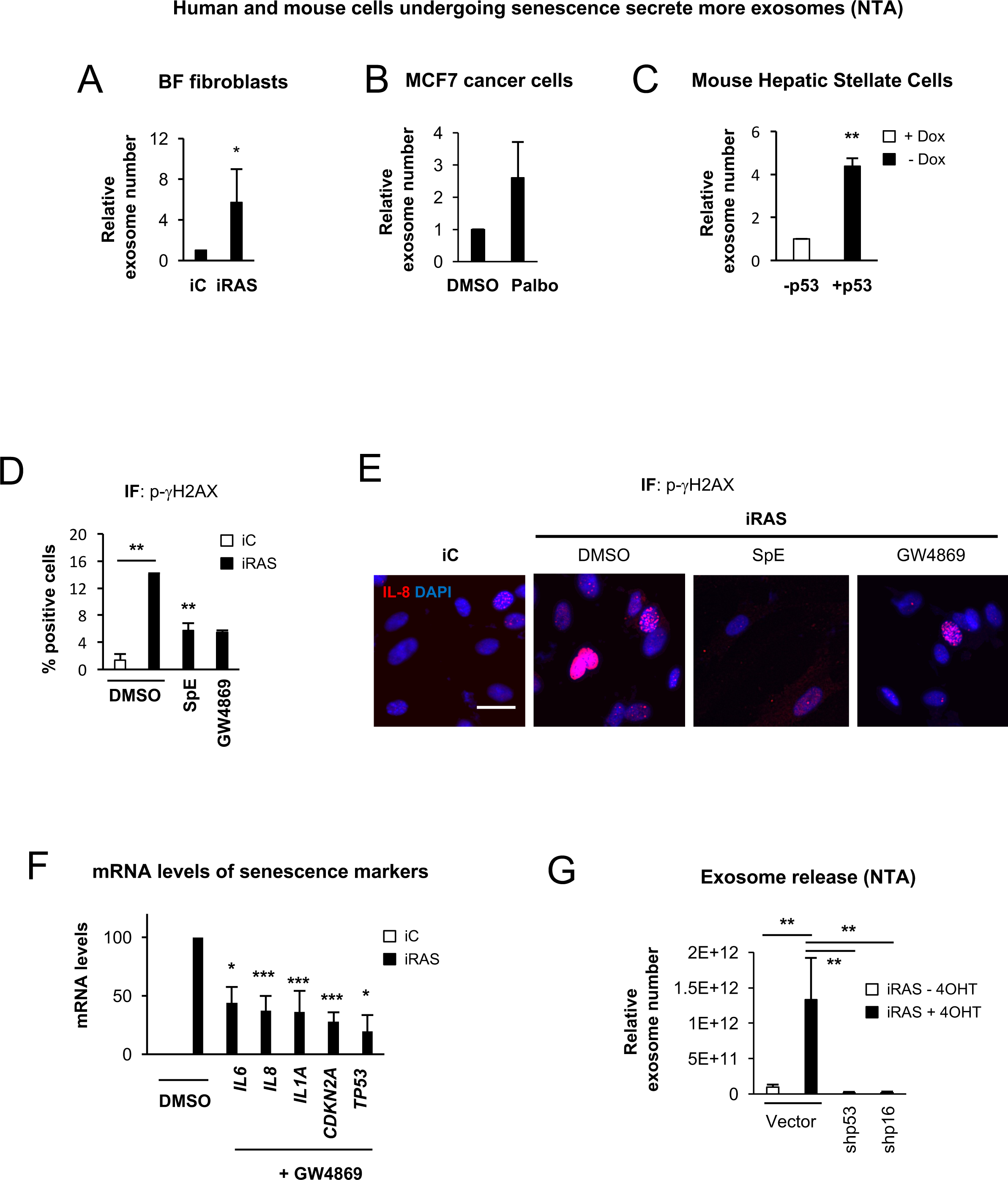

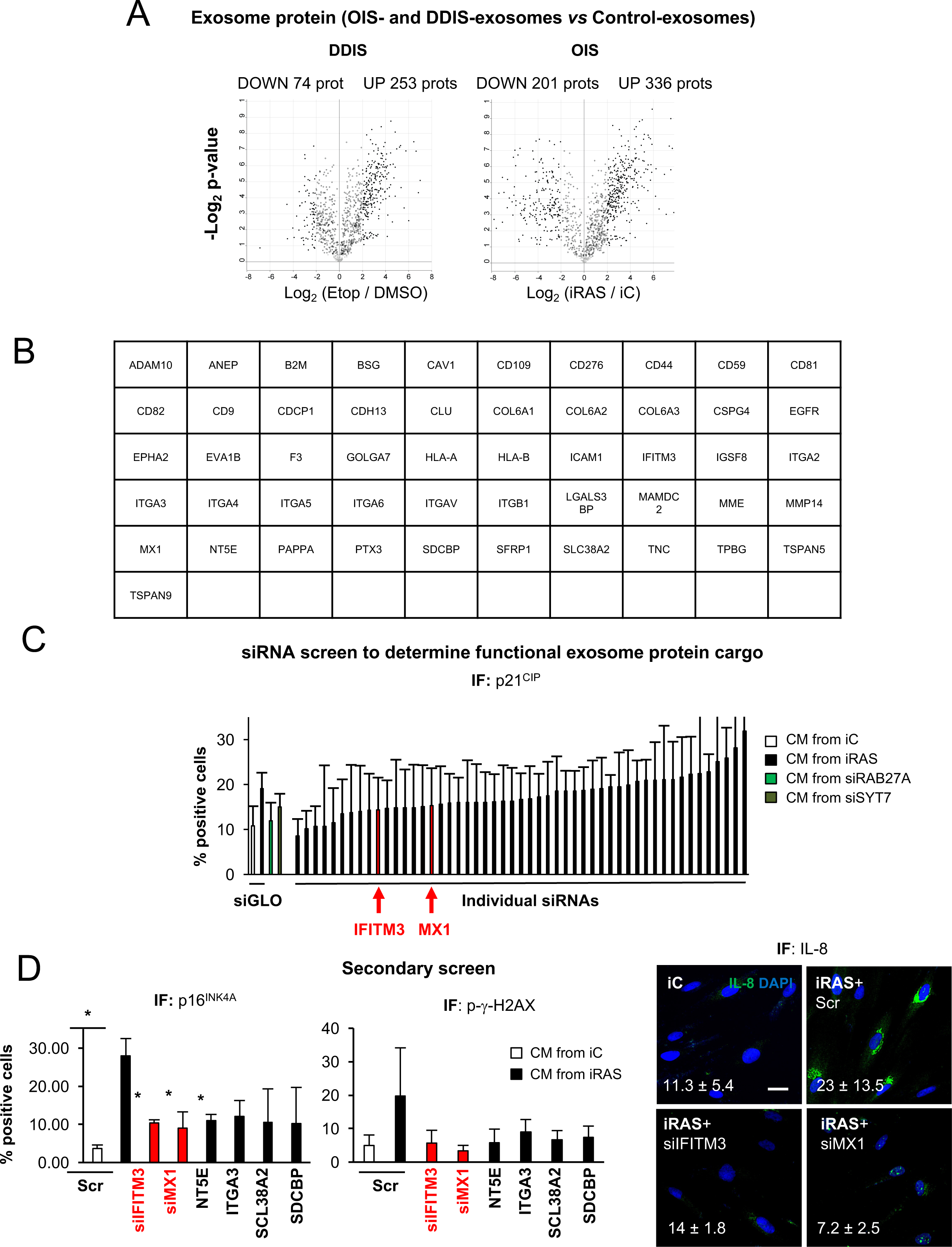

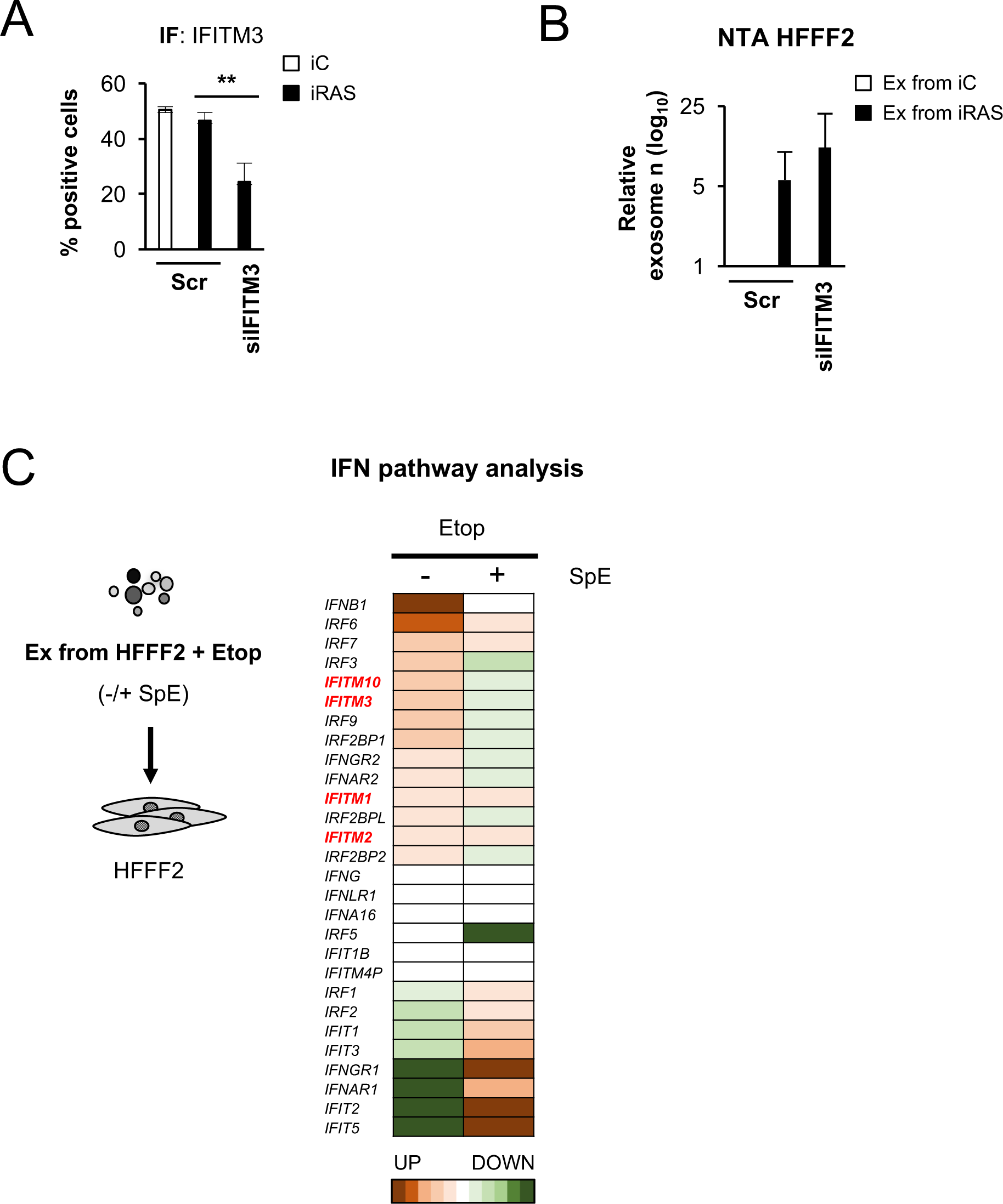

